# Patterns of landscape partitioning indicate low levels of resource competition among neighbouring Guinea baboon (*Papio papio*) parties

**DOI:** 10.1101/2024.09.23.614456

**Authors:** Lisa Ohrndorf, Roger Mundry, Jörg Beckmann, Julia Fischer, Dietmar Zinner

## Abstract

**Background:** Access to critical resources, including food, water, or shelter, significantly determines individual fitness. As these resources are limited in most habitats, animals may employ strategies of landscape partitioning to mitigate the impact of direct resource competition. Territoriality may be regarded as an aggressive form of landscape partitioning but in non-territorial species, other forms exist. Animals living in groups with greater flexibility in their association patterns, such as multilevel societies with fission-fusion dynamics, may adjust their grouping and space use patterns to short-term variations in ecological conditions such as food availability, predation pressure, or the presence of conspecific groups. This flexibility may allow them to balance the costs of competition while reaping the benefits of better predator detection and defence.

**Methods:** We explored patterns of landscape partitioning among neighbouring Guinea baboon (*Papio papio*) parties in the Niokolo-Koba National Park, Senegal. Guinea baboons live in a multilevel society in which parties predictably form higher-level associations (“gangs”). We used four years of locational data from individuals equipped with GPS collars to estimate annual home ranges, home range overlap, and minimum distances between parties. We examined whether food availability and predator presence levels affected the cohesion between parties in 2022.

**Results:** We found substantial home range and core area overlap among parties (33 to 100%). We found no effect of food availability or predator presence on the distance to the closest neighbouring party; the minimum distance between parties was less than 100 m on average.

**Conclusions:** Our results suggest a low level of feeding competition between our study parties. Whether this is a general feature of Guinea baboons or particular to the situation in the Niokolo-Koba National Park remains to be investigated.

## Background

Access to critical resources such as food, water, shelter, or mates can be a significant determinant of the fitness of animals. In habitats where specific resources are limited, there is a high potential for resource competition [1]. An extensive body of theoretical and empirical research targets whether and how animals, particularly primates, adjust aspects of their social organisation, including group size, composition and cohesion, to prevailing ecological conditions in their home ranges [2–10]. These studies have highlighted the importance of resource abundance and distribution in shaping primates’ spatial distribution and social organisation. Specifically, more abundant and evenly distributed resources are expected to result in larger, more cohesive social groups [2,5,11]. Many primate species have been shown to adjust their group size to the distribution and size of feeding patches with smaller feeding parties when feeding patches are small and spatially or temporally clumped, likely resulting in reduced costs of intragroup feeding competition [e.g., 12,13]. While competition for resources presumably sets an upper limit to group size and cohesion in group-living animals, predation pressure is often considered to promote the formation of larger groups. Larger groups can benefit from enhanced predator detection, risk dilution, communal defence, or mobbing of predators [3].

In habitats where ecological conditions are not stable but vary temporally (e.g., seasonally), fission-fusion dynamics can serve as a strategy to balance the costs and benefits of group living [6]. Animals that exhibit fission-fusion dynamics are thought to flexibly adjust their group size, cohesion and composition to different degrees in response to varying ecological conditions [5,6,14]. One way such fission-fusion dynamics can manifest is by partitioning a shared landscape.

Animals may exhibit purely spatial partitioning, where certain parts of the landscape areas are used exclusively. If these areas are defended, this leads to territoriality [5]. In non-territorial species, animals may avoid regions or parts of their home ranges recently used by neighbouring groups or sub-groups in species with fission-fusion dynamics, thus preventing associated costs of aggressive encounters [15–19]. Another way to mitigate resource competition with neighbouring groups that inhabit the same space is to share the same area but temporally avoid other groups or subgroups. Such temporal avoidance could greatly reduce the potential for aggressive encounters and contest competition for valuable resources [18,19].

In this study, we investigated patterns of landscape partitioning among neighbouring Guinea baboon (*Papio papio*) parties in the vicinity of Simenti, Niokolo Koba National Park (PNNK), Senegal. Guinea baboons are group-living, non-territorial primates that are highly spatially tolerant, with most inter-group encounters being neutral or even affiliative [20,21]. They form large multi-male, multi-female groups of 20 to more than 300 individuals. Guinea baboons have a nested multi-level social organisation with fission-fusion dynamics [20]. At the basis of the multi-level society is the one-male unit, consisting of one primary male, one to several females, their dependent offspring, and sometimes secondary males [22]. Several one-male units form parties, which, together with other parties, form gangs [23]. Over 90% of the offspring in a unit are sired by the primary male of a unit at the time of conception, significantly reducing the potential for mate competition [24]. Events of fissions and fusions in Guinea baboons usually happen along consistent lines and most often at the level of parties. Parties can vary in size but frequently consist of more than 20 individuals, including juveniles. The home ranges of parties in the Niokolo Koba National Park in Senegal vary between 20 and 50 km^2^ [21,25]. Like other baboon species, Guinea baboons are considered eclectic omnivores feeding primarily on fruit but also on other plant parts, insects, and occasionally on small birds or mammals [21,25]. In our study area, the most important predators of Guinea baboons are African lions (*Panthera leo*), leopards (*P. pardus*), and spotted hyenas (*Crocuta crocuta*).

We aimed to identify whether and how neighbouring Guinea baboon parties partition a shared space among each other, both purely spatially as well as temporally, and to assess whether this partitioning reflects avoidance or attraction-based patterns. We hypothesize that if neighbouring parties experienced strong resource competition, we would observe *avoidance-based patterns*. This would result in minimal overlap between the core areas of different parties, which could act as a strategy to maintain exclusive access to important resources. Alternatively, parties might avoid each other temporally, maintaining large distances over time to reduce direct encounters while occupying the same overall area.

In contrast, if resource competition between parties is low, we might observe *attraction-based patterns*. In this case, parties would show more overlap in their core areas or maintain closer proximity over time, suggesting that the benefits of group living or the lack of significant competition outweigh the potential costs.

We investigated whether these patterns varied alongside prevailing ecological conditions, namely food availability and predator presence. We anticipated that avoidance-based patterns would become more pronounced when resource availability was low, potentially irrespective of predator presence. Conversely, we expected to see more attraction-based patterns when predator presence was high, as closer cohesion could provide protection from predators.

## Methods

### Study site

The data collection for this study took place at the long-term field site of the Centre de Recherche de Primatologie (CRP), Simenti (13°01‘34‘‘ N, 13°17‘41‘‘ W) in the Niokolo-Koba National Park (PNNK) in south-east Senegal (Fig. 1). The study site is part of the Sudanian and Sahelo-Sudanian climatic zone with pronounced seasonality and high seasonal variability in rainfall [26]. The average annual precipitation in Simenti is around 950 mm. The rainy season lasts from June to October, with May and mid-October constituting transitional periods with minor and variable rainfall [26]. The vegetation represents a mosaic of grasslands, wooded savannahs, and gallery forests along streams and other perennial water bodies [26,27]. Several habituated Guinea baboon parties live at the study site, followed extensively by researchers since 2007.

**Fig. 1:**
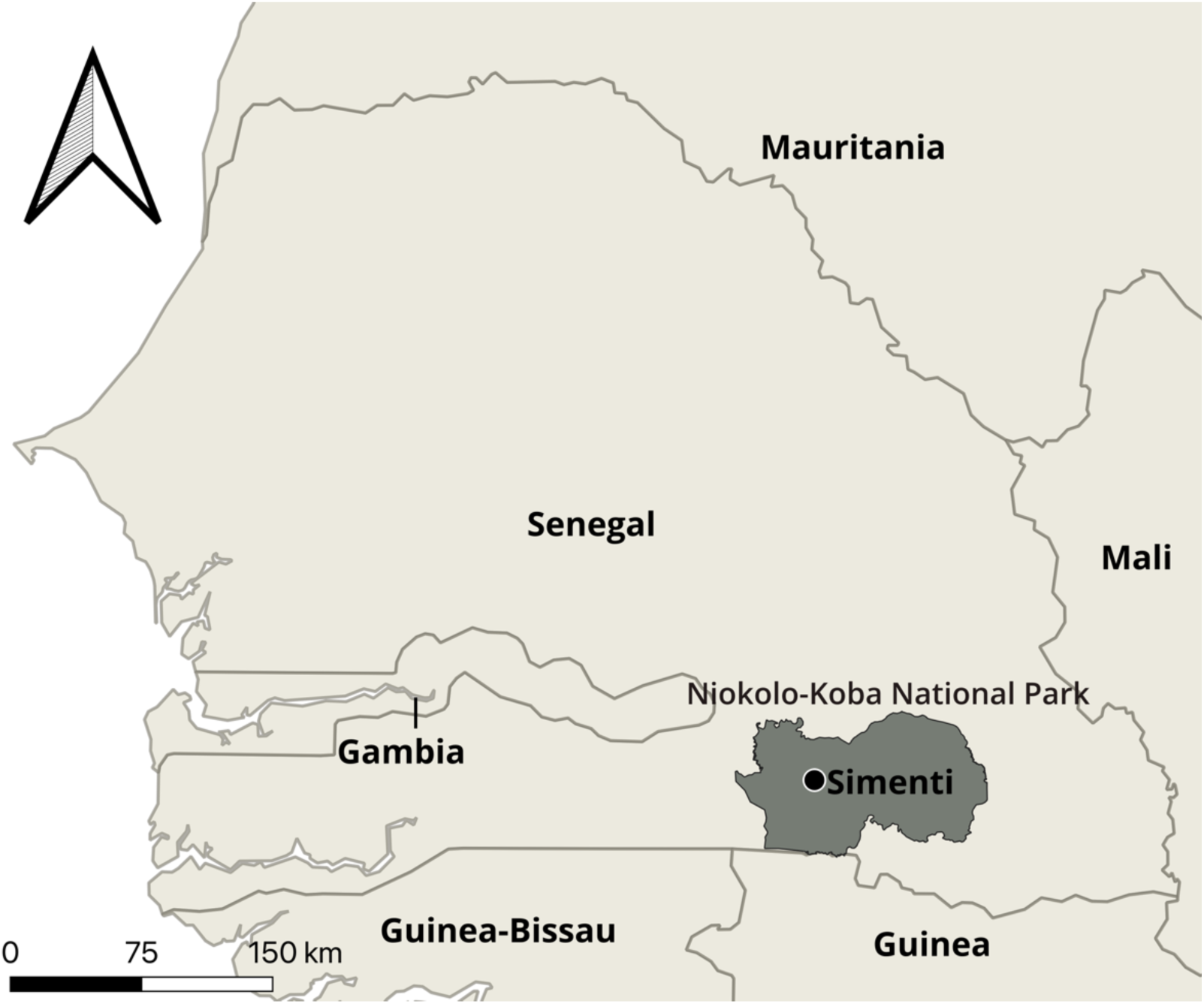
Location of the field site in Simenti (black dot) within the Niokolo-Koba National Park.

### Data collection

#### GPS Data

To assess the spatial distribution of Guinea baboon parties across the study area, we equipped eight adult males, each representing the location of their entire party (P5, P6I, P6W, P7, P9B, P13, P15, P17), with GPS collars (Tellus 2 Basic Light) with integrated drop-off mechanisms from January to December 2022. Party membership was assigned based on spatial proximity, and individuals spent most of their time with members of their parties [20]. The GPS collars were programmed to take locational fixes every two hours during the day (06:00 to 18:00) and an additional three fixes during the night to mark the location of the sleeping site (21:00, 00:00, 03:00). We downloaded GPS data from the collars using a UHF antenna monthly.

In addition to the data from 2022, locational data from a previous study were available for 2010 to 2012 [20]. This dataset comprised locational data with identical sampling intervals for eleven individuals (four females) from six different parties in 2010, seven individuals (two females) from six parties in 2011 and five individuals (one female) from five parties in 2012. As this dataset also contained individuals from unhabituated parties, we excluded these from further analyses to remove any effects of habituation as a potential confounding factor.

#### Capture and collaring

Collaring for this study took place on five days between January 16 and January 21, 2022. We located the targeted baboons and their parties in the morning, followed them and short-term immobilised them covertly (without either the target animals themselves or surrounding individuals seeing the shot) while they were ranging with their parties by darting via a blowpipe (all darting equipment was from TeleDart: blowpipe B16, calibre 16 mm, length 160 cm, 2 ml darts (BD2) with plastic stabilisers (BST16) and plain needles 1.5 x 38 mm [TDN1538LL]). The initial dose for chemical immobilisation per baboon was 100 mg ketamine, 20 mg xylazine, and 2 mg atropine. As soon as the target baboon was fully immobilised and all other baboons had left the area (at least 300 m from the immobilized target baboon), we moved the baboon to a well-shaded area, blindfolded the male and started to monitor vital parameters (oxygen saturation, heart rate, and internal body temperature). We weighed the animal using a spring scale. The mean body mass of males was 21.5 kg ± 1.9 SD (range: 19-25 kg, n=8). After collaring, we moved the animal close to a small tree in the shade and administered 1.5 mg of atipamezole as an antidote against xylazine. We closely surveyed the baboon from a distance until it was fully awake and able to move safely. All collared baboons successfully rejoined their respective parties on the same or the following day. A veterinarian of the Senegalese Direction des Parcs Nationaux (DPN), and a representative of the veterinary service of Tambacounda accompanied and supervised the entire capture and collaring procedure. The collaring was approved by the Ethics committee of the German Primate Center (document number E4-21). For the collaring procedures from 2010 to 2012, see Knauf et al. [28].

#### Phenological data

To estimate food availability across the home range of our study parties, we recorded the phenological state of known feeding tree species from November 2021 to March 2023. To this end, we selected 31 feeding tree species that our study parties fed on regularly during a previous study [21] (Table S1). For each of these feeding tree species, we selected ten individual trees on average (range 1-13) that, if possible, were distributed evenly across the different habitat types our study population occupies. The trees were marked with aluminium tags and visited at the beginning of each month. We recorded each tree’s phenological state according to its phenological activity (eight levels: none, young leaves, flower buds, flowers, young fruits, intermediate fruits, ripe fruits, mature leaves). We considered trees bearing ripe or intermediate fruit as “providing food”. As our study population also consumed the flowers of certain tree species, we included months in which those trees were flowering as “providing food”. We then calculated the food availability from the proportion of trees providing food per species and divided them by the total number of tree species observed.

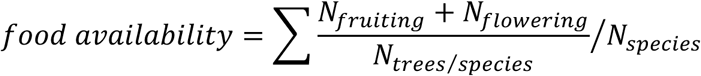

#### Predator presence

To estimate the predator presence at our study site, we used non-baited motion-triggered camera traps distributed across 37 km^2^, covering most of our study parties’ home ranges. The cameras were deployed on a grid of 1 x 1 km, with roughly one camera per km^2^ (Fig. 2).

**Fig. 2:**
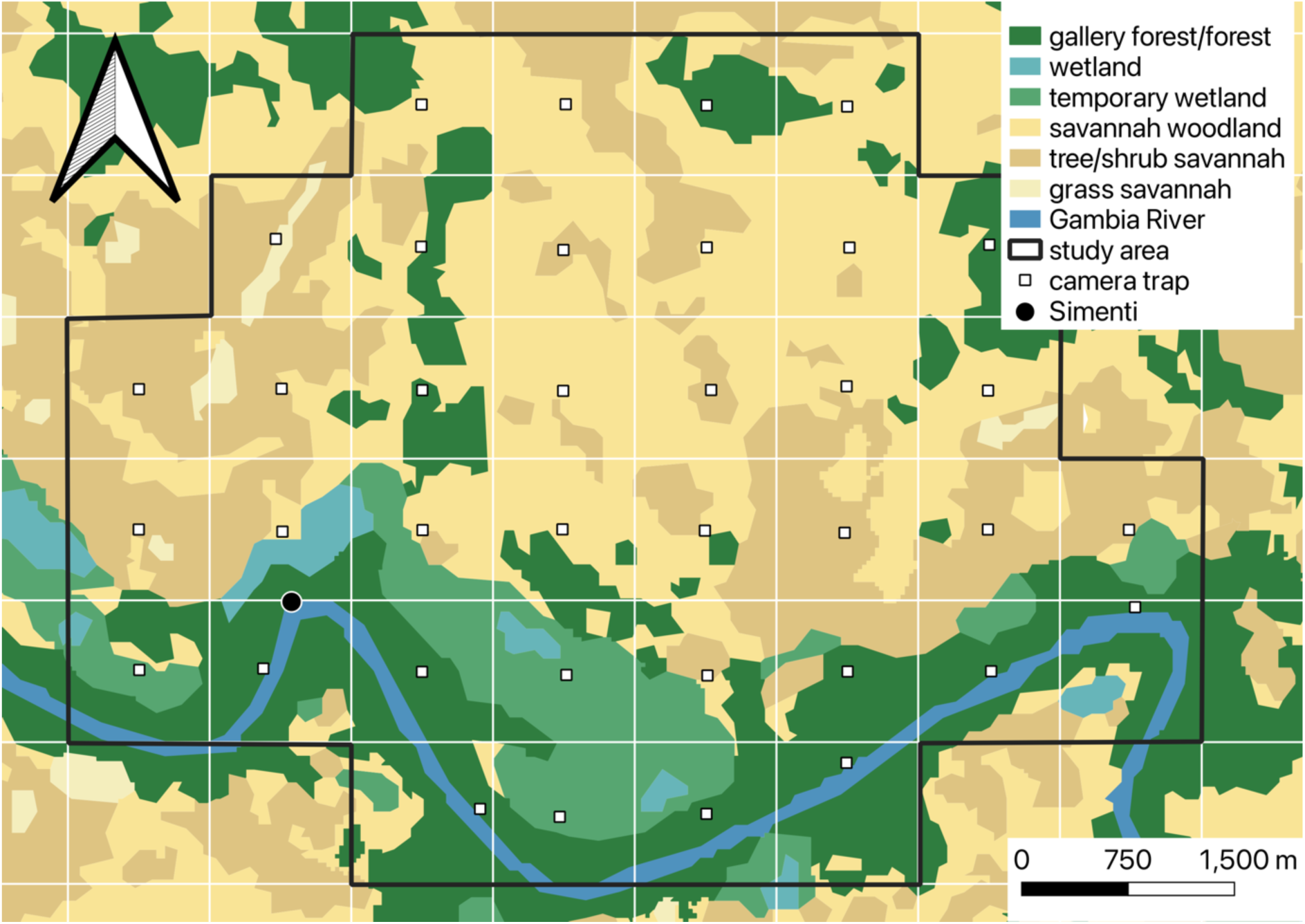
Distribution of camera traps and habitat types within the study area. White lines represent 1 km x 1 km grid cells. The location of the field site of the CRP Simenti is indicated by a black dot and the positions of the camera traps are depicted as white squares.

All cameras except one were installed facing towards frequently used animal trails. We installed cameras in February 2022 and retrieved them in March 2023. In the meantime, we exchanged batteries and SD cards monthly. During the rainy season, we cleared an area of ca. 5 m around each camera from grass to ensure unobstructed visibility and avoid excessive triggering of the cameras by vegetation. The cameras were programmed to take three consecutive pictures upon being triggered, with a 1-min interval between triggers. The imagery obtained from this camera-trapping grid was annotated using the online platform *Agouti* [29]. The AI-assisted annotation was manually checked for all images. We recorded animal species and the number of individuals for all images taken by the camera traps. We considered all sightings of lions, leopards, or hyenas within the camera-trapping grid as a predator encounter. In addition to data from the camera trapping grid, we used ad libitum data on all signs of predators (tracks, scat, calls, sightings) recorded at the field site during the study period. From both camera trapping data and ad libitum observational data, we counted the number of predator encounters in the study area at 14-day intervals. As we could not be sure about the most relevant time window for assessing perceived predation pressure in Guinea baboons, we also evaluated predator encounters over shorter (2 and 7 days) and longer (30 days) intervals.

### Statistical analyses

#### Spatial landscape partitioning

We estimated each party’s annual home ranges and core areas using the package *amt* [30] in R version 4.3.1 [31]. We delineated home ranges and core areas as the 95% and 50% contour levels obtained from Kernel Density Estimation (KDE), respectively. We used a rule-based ad hoc approach (SCALEDh) to select bandwidths within a search range of 0.01 and 1 of REFh [32]. For 2010-2012, we calculated home ranges and core areas for all individuals that belonged to one party collectively, as several individuals per party were collared. Further, as there was substantial overlap between collaring periods in 2011 and 2012, we pooled locational data for individuals from the same parties for overlapping periods. We then used the function ‘hr_overlap’ from the package *amt* [30] to calculate home range overlap between parties per year.

#### Temporal landscape partitioning

To assess temporal patterns of cohesion between our study parties, we calculated the average distance between pairs of collared individuals, each representing their entire party’s location, for each day they were observed together. Then, we identified the shortest distance between all dyads of parties until each party was represented at least once for every day it was observed on the same day as another party. This procedure allowed us to determine the minimum distance between neighbouring parties each day (Fig. 3).

**Fig. 3:**
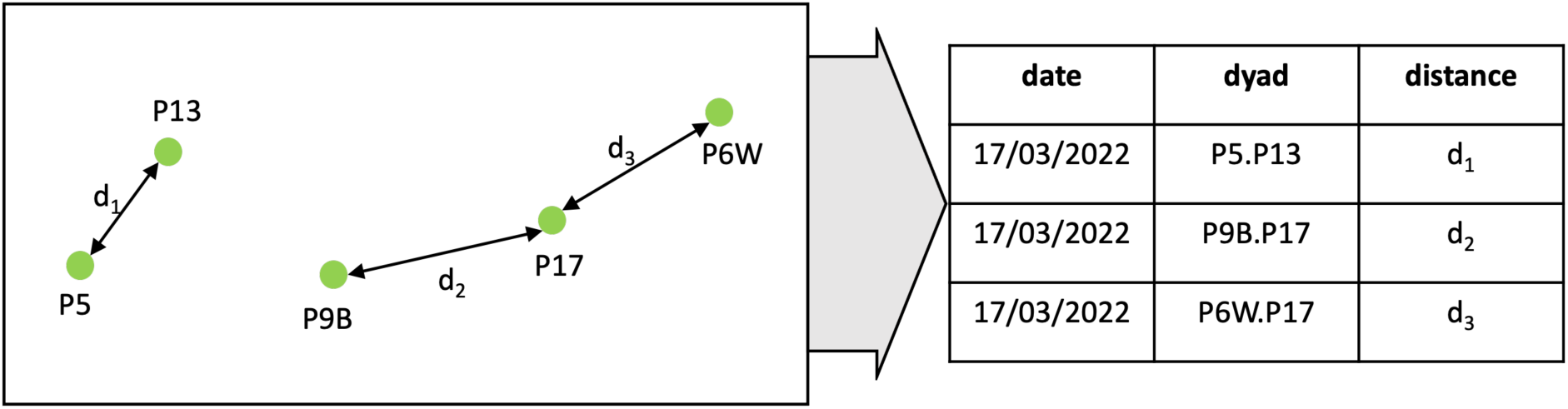
Exemplary depiction of the assessment of minimum distances between parties. For the exemplary day, GPS data were available for five parties (P5, P6W, P9B, P13, P17). We averaged the distances between parties across the day. We then identified the closest neighbouring dyads (P5 and P13, P9B and P17, P6W and P17) and the Euclidean distances between them (d_1_-d_3_).

We used minimum rather than average distances between parties because, in a limited space, movements away from one party would often lead to proximity to another, leading to averages not being able to capture the spatial relationships we were interested in.

We also used this approach to validate our assumption that one individual of a party is representative of the entire party’s location. To this end, we compared minimum distances between individuals from the same party to minimum distances between members of different parties from 2010 to 2012 (Fig. S1).

#### Modelling

To assess whether patterns of temporal landscape partitioning varied with food availability and predator presence, we used data on minimum distances between parties, food availability, and predator presence only from 2022. We fitted a multiple membership model using the function *brm* of the package *brms* version 2.20.4 [33] in R version 4.3.1 [31]. We log-transformed the minimum distance between parties to increase the probability of model convergence. We included log distance as the response variable, the food availability score, and the number of predator encounters within 14 days prior to the observation as predictors. We included the IDs of the two individuals of a dyad (as a multi-membership term) and also dyad ID as random intercepts effect and random slopes of food availability and predator presence within 14 days in the model to avoid overconfident model estimates [34,35]. A multi-membership term means that only one random intercepts effect is estimated for both individuals involved in a dyad. This accounts for the possibility that some parties are generally closer to others than other parties. The dyad effect in term accounts for the possibility that some parties tend to associate with one another more than others. We fitted a multi-membership model since the two individuals of a dyad could not be unambiguously assigned to two different random effects variables. We checked for sufficient variation in food availability and predator presence within each individual before fitting the model. We also included parameters for the correlations among random intercepts and random slopes in the model. As we received a warning about divergent transitions during warm-up with the default settings of *brm*, we set adapt_delta to 0.9.

We fitted three additional models, including the number of predators per 2, 7 and 30 days instead of 14 days as predictors, but with an otherwise identical structure to ensure model results were not biased by an inappropriate choice of the time window used to assess predator presence. To check whether the resulting patterns were solely driven by Guinea baboon parties aggregating at sleeping sites in the mornings or evenings, we fitted four additional models with identical structures, using only the minimum distance between parties at noon as the response.

## Results

Across all study years, we obtained a total of 58,404 locational fixes during the day, excluding those taken during the night (21:00, 00:00, 03:00). 17365 of these locational fixes were collected in 2022 during 347 unique days. The GPS collars lasted 316 days on average, with one collar (for P6I) failing after 162 days due to water damage.

### Spatial patterns of landscape partitioning

In 2022, our study parties consisted of 25 individuals on average (including males, females, and juveniles), ranging from seven individuals in P15 to 41 individuals in P5. We collectively estimated annual home ranges and core areas for parties P4, P5, P6, and P9 for 2010, as well as for 2011 to 2012. For 2022, we estimated the home ranges and core areas of parties P5, P6I, P6W, P7, P9B, P13, P15, and P17. The average home range size in 2022 was 38.4 km^2^ (range 20.6-45.9 km^2^), and core areas were around 8.5 km^2^ (range 3.5-10.9 km^2^). The average overlap between home ranges of parties in 2022 was 89% (range 45-100%, Table 1). Core area overlap between parties in 2022 was 80% on average (range 33-98%, Table 2).

**Table 1:** Overlap between home ranges (KDE, 95% contour level) of parties observed in 2022.

|  | P5 | P6I | P6W | P7 | P9B | P13 | P15 | P17 |
| --- | --- | --- | --- | --- | --- | --- | --- | --- |
| P5 |  | 0.96 | 0.96 | 0.50 | 0.96 | 0.99 | 0.98 | 0.96 |
| P6I | 0.96 |  | 0.96 | 0.73 | 0.94 | 0.96 | 0.84 | 0.86 |
| P6W | 0.91 | 0.94 |  | 0.45 | 0.98 | 0.87 | 0.92 | 0.87 |
| P7 | 1.00 | 0.99 | 1.00 |  | 1.00 | 1.00 | 1.00 | 1.00 |
| P9B | 0.92 | 0.95 | 0.99 | 0.46 |  | 0.89 | 0.91 | 0.88 |
| P13 | 0.99 | 0.96 | 0.98 | 0.50 | 0.98 |  | 0.98 | 0.92 |
| P15 | 0.99 | 0.98 | 0.96 | 0.51 | 0.97 | 0.99 |  | 0.97 |
| P17 | 0.82 | 0.97 | 0.90 | 0.45 | 0.89 | 0.84 | 0.82 |  |

Values represent the proportion of overlap (0 = no overlap, 1= complete overlap).

**Table 2:** Overlap between core areas (KDE, 50% contour level) of parties observed in 2022.

|  | P5 | P6I | P6W | P7 | P9B | P13 | P15 | P17 |
| --- | --- | --- | --- | --- | --- | --- | --- | --- |
| P5 |  | 0.77 | 0.95 | 0.40 | 0.94 | 0.97 | 0.95 | 0.96 |
| P6I | 0.75 |  | 0.68 | 0.46 | 0.65 | 0.75 | 0.53 | 0.59 |
| P6W | 0.87 | 0.71 |  | 0.33 | 0.93 | 0.79 | 0.85 | 0.79 |
| P7 | 0.97 | 0.76 | 0.98 |  | 0.97 | 0.96 | 0.96 | 0.98 |
| P9B | 0.87 | 0.73 | 0.97 | 0.33 |  | 0.81 | 0.84 | 0.82 |
| P13 | 0.96 | 0.79 | 0.95 | 0.37 | 0.93 |  | 0.93 | 0.89 |
| P15 | 0.98 | 0.87 | 0.93 | 0.41 | 0.93 | 0.98 |  | 0.96 |
| P17 | 0.84 | 0.78 | 0.92 | 0.36 | 0.92 | 0.88 | 0.80 |  |

Values represent the proportion of overlap (0 = no overlap, 1= complete overlap).

### Food availability

Food availability scores ranged from 0.03 to 0.33 across the study period (Fig. 4A). February, March, and April 2022 had the highest scores. In contrast, June, July, and August had the lowest scores.

**Fig. 4:**
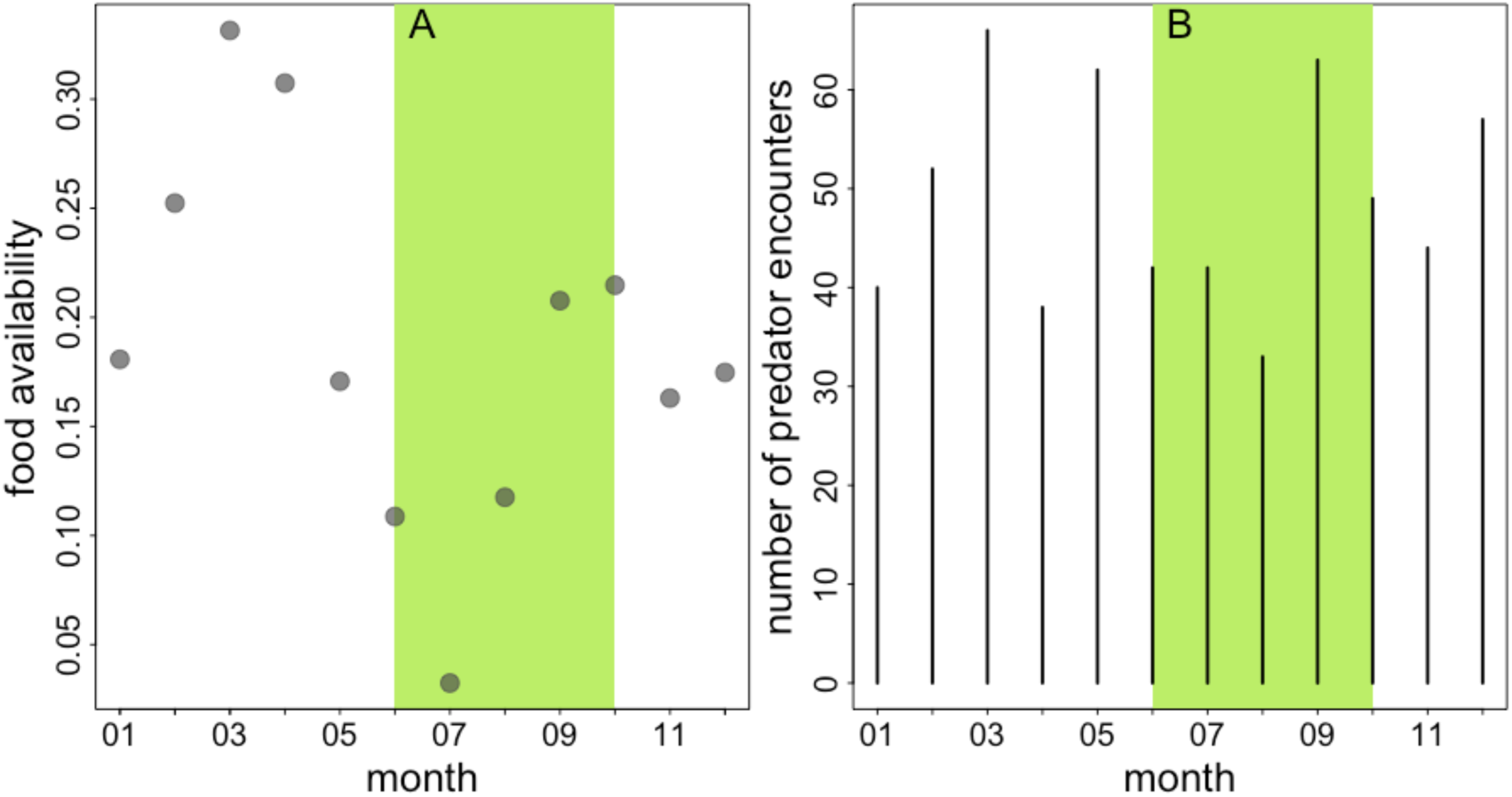
Monthly food availability scores (A) and number of predator encounters per month (B) in 2022. The light green area indicates the rainy season (June-October).

### Predator presence

In 2022, we registered 588 predator encounters, 376 of which were ad libitum records and 212 were camera trap images of predators. Most records were from spotted hyenas (254), followed by lions (211), and leopards (118). Of the remaining five records, one was a sighting of a pack of eleven African wild dogs (*Lycaon pictus*), and four images could not be reliably identified as either leopard or lion.

For the 14-day interval, there were 25 predator encounters on average (median, range 17 – 42). For the 7-day interval, the average number of predator encounters was 13 (median, range 7 – 27). The average number of predator encounters for the 30-day interval was 53 (median, range 39 – 82), and for the 2-day interval, it was around 3 (median, range 2-13).

### Temporal patterns of landscape partitioning

The average minimum distance between neighbouring Guinea baboon parties in 2022 was 81.4 m (median, 47.1 – 207.3 m IQR) (Fig. 5). Throughout the year, we observed short minimum distances between neighbouring parties (Fig. 6). There were many instances of close proximity and fewer instances of larger distances between parties. However, in April and May, these minimum distances were notably larger, increasing from approximately 50 m to 100 m.

**Fig. 5:**
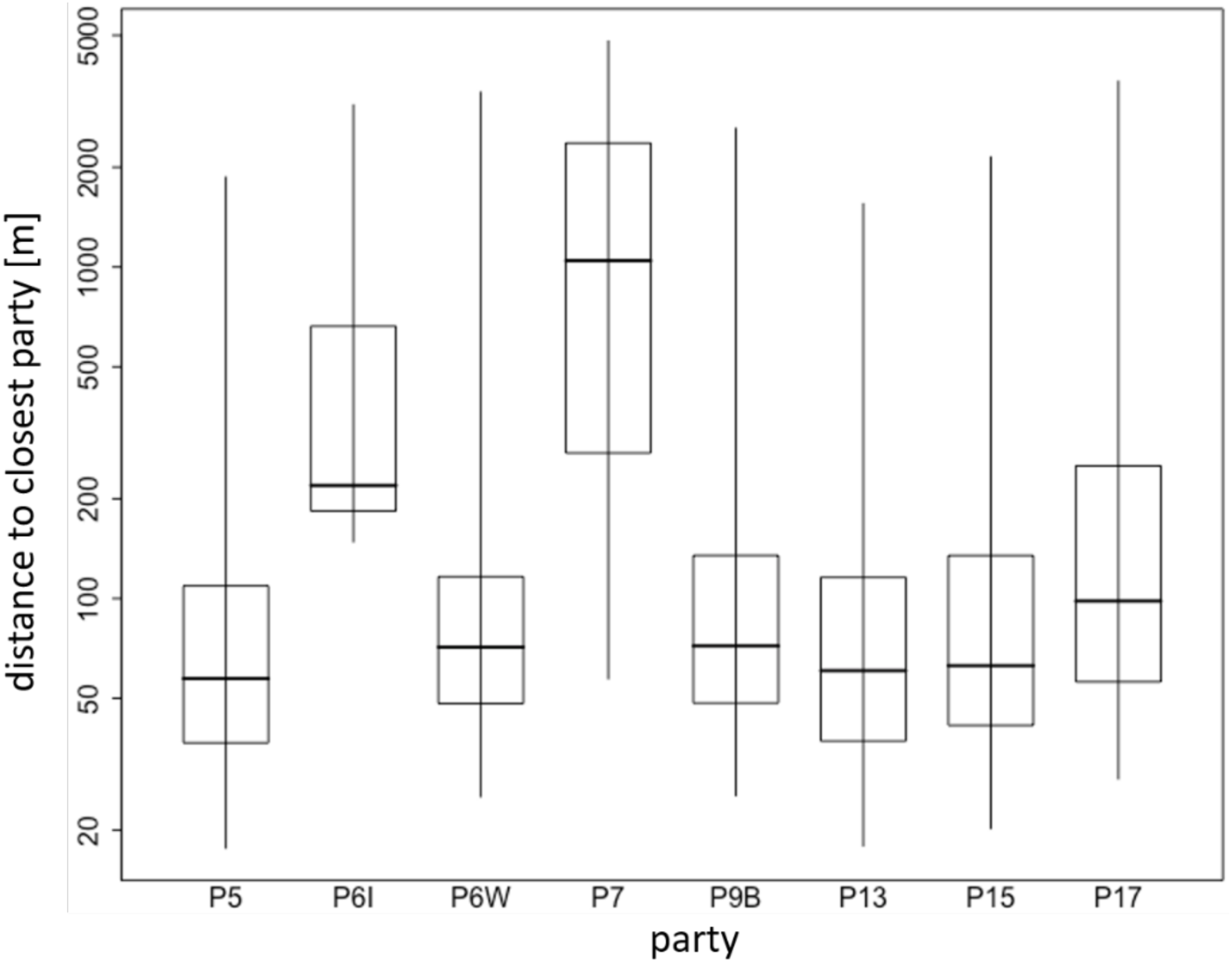
Temporal cohesion between parties in 2022. Boxplots depict the median (black line) and IQR with the lower (25%) and upper (75%) quartile. Whiskers represent the 2.5^th^ and 97.5^th^ percentiles. The distance to the closest party (y-axis) is depicted on a log-scale for visual clarity.

**Fig. 6:**
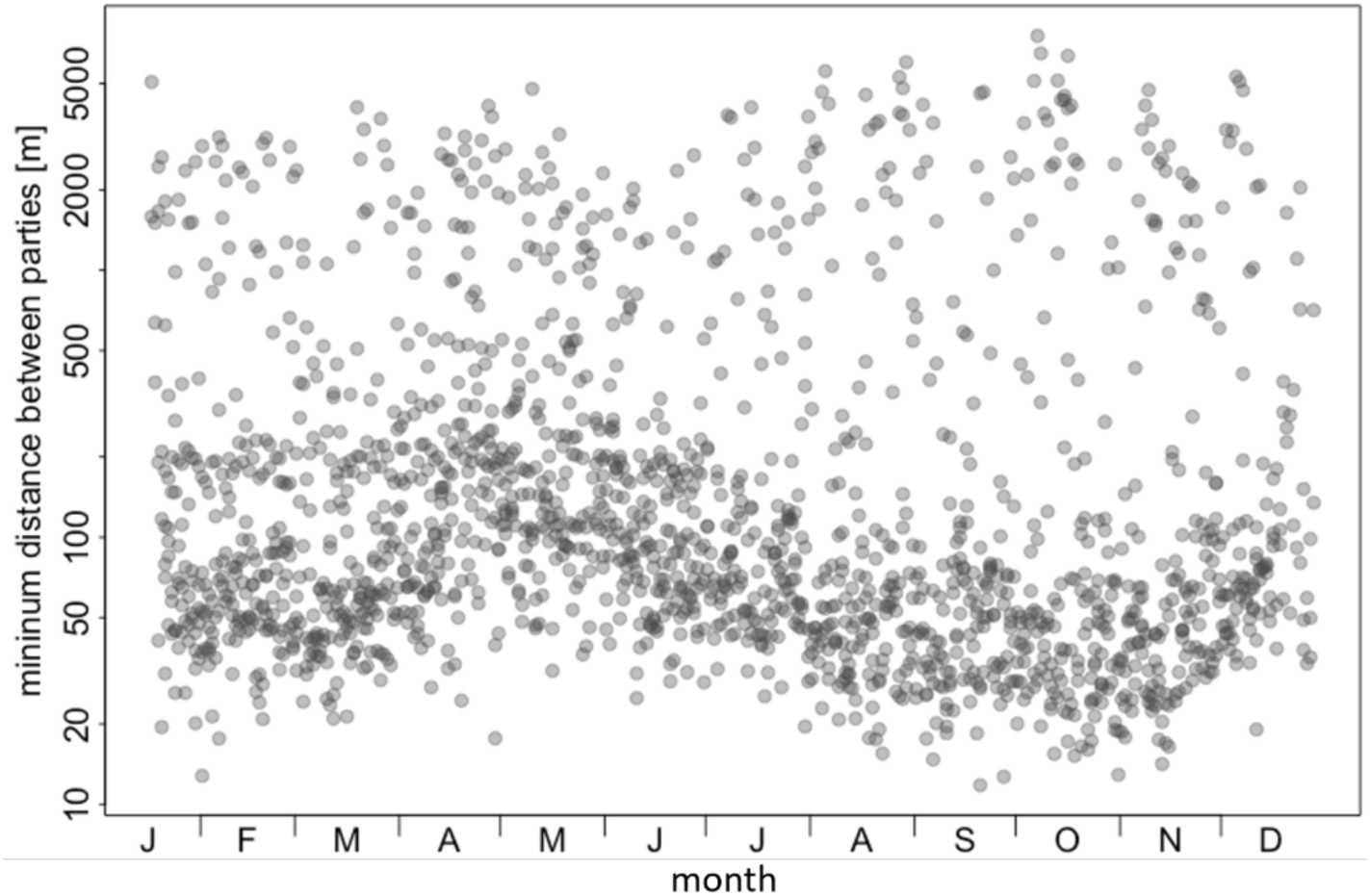
Minimum distances between parties throughout 2022. The minimum distance between parties (y-axis) is depicted on a log-scale for visual clarity.

We did not find evidence for our study parties to adjust their space use patterns alongside varying levels of food availability (Fig. 7A) or predator presence (Fig. 7B; Table 3). The estimated effect sizes were marginal, regardless of the time interval chosen when calculating predator presence (Tables S5-S7). Model results also did not reveal any effect of predator presence and food availability when considering only waypoints taken at noon (Tables S8-S11).

**Fig. 7:**
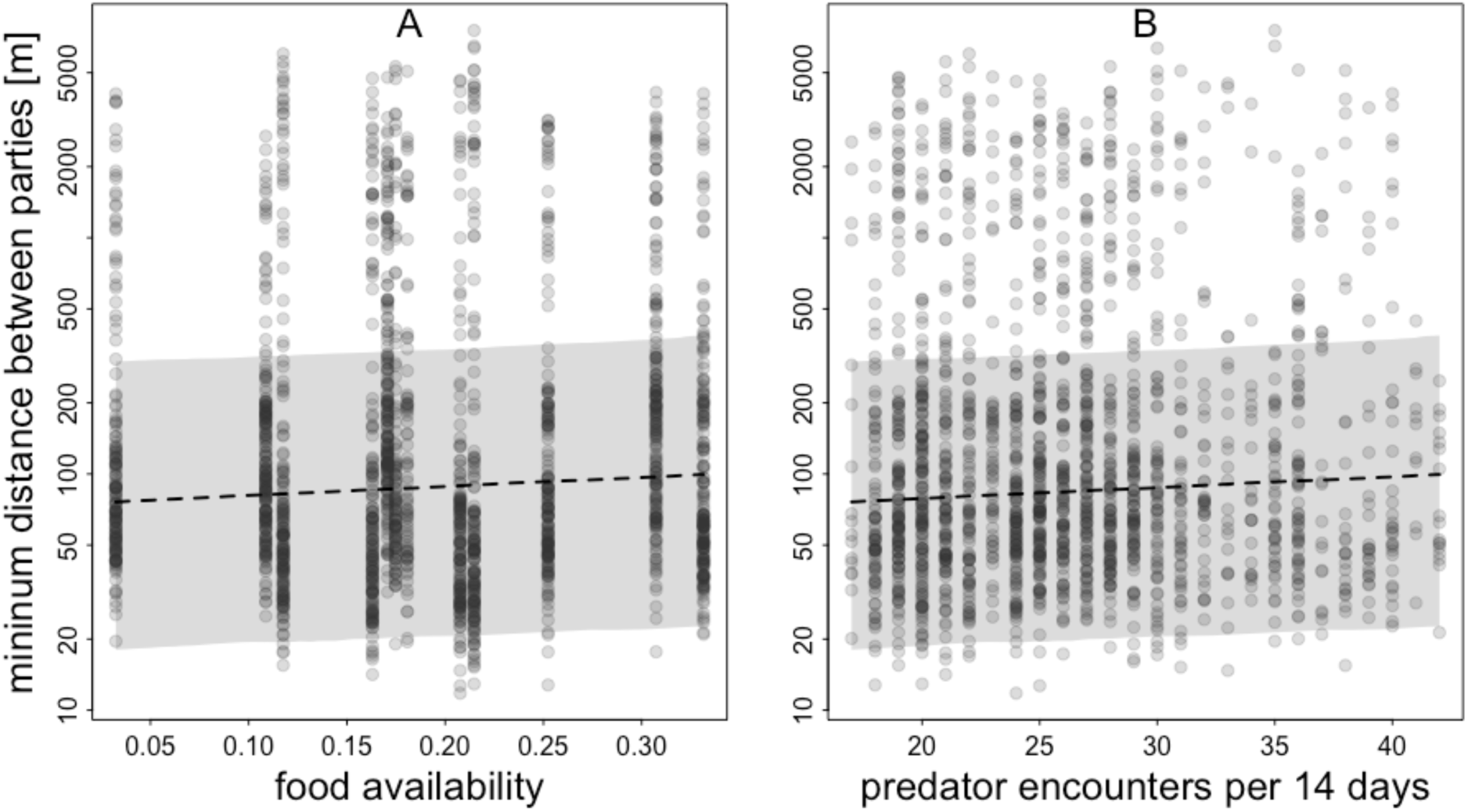
Model results for the effect of food availability (A) and the number of predator encounters per two weeks (B) on the minimum distance between parties in 2022. For (A) predator presence was centred to a mean of zero and for (B) food availability was centred to a mean of zero. The minimum distance between parties (y-axis) is depicted on a log-scale for visual clarity. The fitted mean is depicted as a dashed line, and confidence intervals are shaded in grey.

**Fig. 8:**
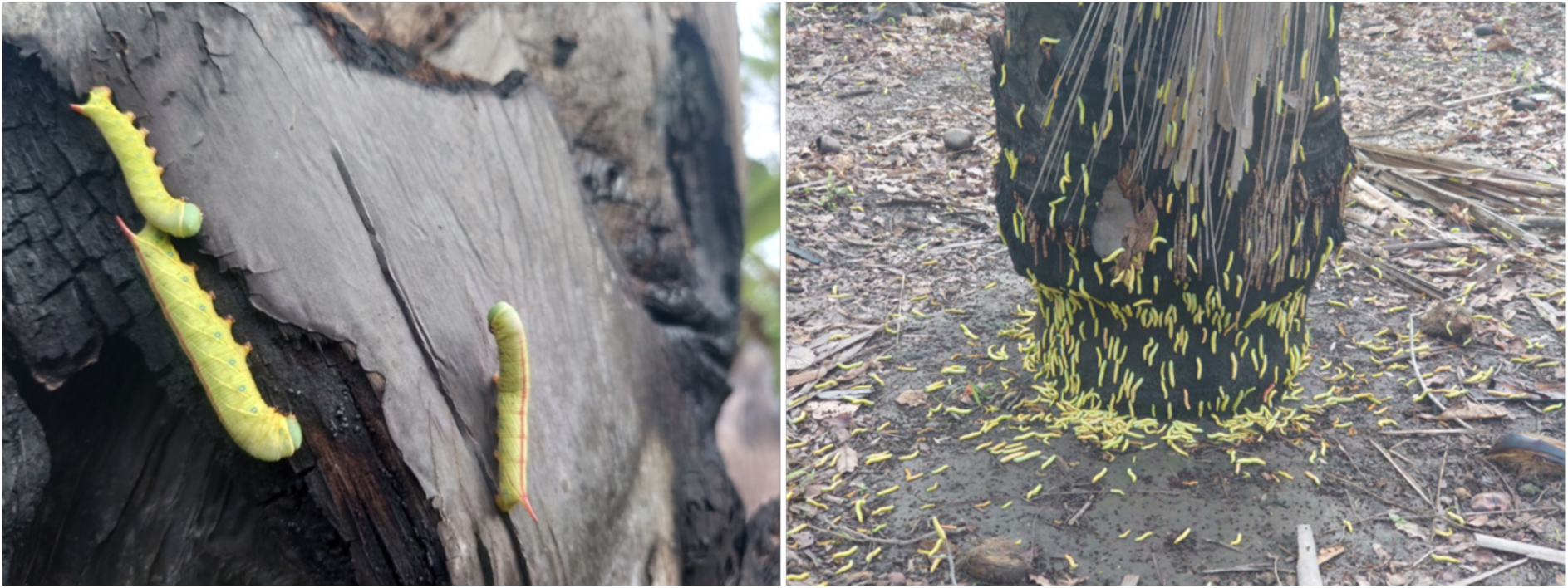
Caterpillars of a species of Sphingidae during a mass occurrence in 2022. Pictures by Marc Mönich.

**Table 3:** Model results on spatiotemporal landscape partitioning between neighbouring parties in response to food availability and number of predator encounters within two weeks (estimates, standard errors, credible intervals, Rhat, as well as Bulk and Tail Effective Sample Sizes).

| Term | Estimate | Est.Error | Cl<sub>lower</sub> | Cl<sub>upper</sub> | Rhat | Bulk_ESS | Tail_ESS |
| --- | --- | --- | --- | --- | --- | --- | --- |
| Intercept | 5.14 | 0.75 | 3.64 | 6.63 | 1.00 | 866 | 1440 |
| pred.enc.14 | 0.00 | 0.01 | -0.02 | 0.02 | 1.00 | 1418 | 1836 |
| food.score | 0.13 | 0.52 | -0.86 | 1.20 | 1.00 | 2424 | 2750 |

## Discussion

### Spatial landscape partitioning

Our analyses revealed substantial overlap between home ranges and core areas of most of the observed Guinea baboon parties. Almost all parties’ home ranges overlapped by 90%, and core areas overlapped by 80% on average, except for one party (P7). Although most individuals were habituated to the presence of researchers in the field, this party was not part of our regular study parties but probably belonged to a different band than the other study parties. While the home ranges of the other study parties overlapped relatively little with that of P7, P7’s home range was considerably smaller and completely enclosed within the home ranges of the other parties.

As troops in chacma baboons and bands in hamadryas baboons correspond more closely to the level of gangs rather than parties in Guinea baboons [20, 38], our results of home range overlap cannot be compared to other studies directly. However, our results also indicate considerable overlap (45-100%, depending on the perspective of comparison) between neighbouring gangs. The amount of home range overlap between Guinea baboon parties of the same as well as of different gangs in Simenti was considerably higher than what was reported for chacma baboon (*P. chacma*) troops in Suikerbosrand Nature Reserve in South Africa. The average overlap in South Africa was ca. 5% (median, range 0 – 53.72%) during the wet season and ca. 3% (median, range 0 – 45.34%) during the dry season [36]. In Erer-Gota, Ethiopia, home ranges of bands of hamadryas baboons (*P. hamadryas*) overlapped by about 50% with at least seven neighbouring bands [37]. Similar to what we observed in Guinea baboons in Simenti, Altmann & Altmann [39] and Markham et al. [18] found a pronounced overlap between home ranges of neighbouring troops of yellow baboons (*P. cynocephalus*) in Amboseli, Kenya. They concluded that, most likely, there is no part of their study troops’ home ranges that they occupy exclusively. Yellow baboons did, however, spend less time in overlapping areas and had fewer encounters with neighbouring groups than would be expected by chance, suggesting an avoidance-based pattern of landscape partitioning [18].

Similarly, owl monkeys (*Aotus azarae*) in Northern Argentina showed relatively pronounced overlap between home ranges (48% ± 15%) but minimal overlap between core areas of 11% on average, likely to maintain exclusive access to clumped resources in their core areas [40]. A study on space partitioning in mountain gorillas (*Gorilla beringei beringei*) found home range overlap between neighbouring groups of 42% on average (median, range 9.7 - 94.8%), but substantially lower overlap between core areas, with an average of under 10% for 7 out of 10 study groups The authors concluded that this might be a strategy to maintain exclusive access to vital resources located in the core areas for a non-territorial species [19].

Contrary to our initial assumption, the Guinea baboons did not seem to maintain exclusive access to essential resources in the core areas. Instead, they exhibited similarly high levels of overlap in these areas, as seen across their entire home range. We therefore conclude that Guinea baboon parties do not exhibit a purely spatial pattern of landscape partitioning. This may be due to the high population size of Guinea baboons in the Niokolo-Koba National Park. Population estimates in 1998 and 2018 suggested a population size between 100,000 and 250,000 individuals [41, 42], translating to population densities of 10.9 individuals per km^2^ or 27.4 individuals per km^2^, respectively (but see also Sharman [25] with 8.7 baboons/km^2^). The population of Guinea baboons in the Niokolo-Koba National Park may thus already be close to carrying capacity, and all suitable habitat for Guinea baboons is likely occupied, so it may simply not be possible to avoid neighbouring parties purely spatially.

### Temporal landscape partitioning

When looking at the temporal patterns of cohesion between parties, we found that most parties stayed in close spatial proximity to at least one other party, with an average minimum distance of less than 100 m. The exception was again party P7, which seemed further away from the other study parties on average. P7 is not one of our regular study parties and belongs to a different gang; unsurprisingly, we found greater distances between them and the other parties. Given that we equipped only eight parties that inhabit the study area with GPS collars, it is very likely that P7 also stayed near other non-collared parties.

Although we observed an increase in minimum distances between parties from around 50 m to approximately 100 m in April and May, this change likely has no substantial biological implications related to predator detection and defence, or feeding competition between parties. It is possible that this increase reflects a larger group spread while parties remained in close proximity or even mingled with other parties.

Our model results revealed no effect of food availability or predator presence on the cohesion between parties, at least at the spatiotemporal resolution we used to assess food availability and predator presence,. Despite substantial variation in both ecological variables across the study period, Guinea baboons stayed in very close spatial proximity regardless of food availability or predator presence. This observation did not support our hypothesis that Guinea baboons would flexibly adjust their proximity to neighbouring parties to reduce feeding competition when food availability was lower or to increase the potential for predator detection and defence when predator presence was higher.

Other studies on fission-fusion societies have produced mixed results regarding how ecological conditions affect party or subgroup cohesion, both within and among species. For instance, Spider monkeys (*Ateles geoffroyi*) exhibited varying association patterns, affected by seasons, food availability, and precipitation. However, those effects were highly context-dependent, varying with the sex of the study animals, their habitats, and whether association or proximity patterns were considered [43,44]. Similarly, party sizes of chimpanzees (*Pan troglodytes*) have been linked to food availability [12,45], predation pressure [46] and the number of receptive females within the party [47,48] in some studies but not others [49].

While these species exhibit more individualistic patterns of fission-fusion dynamics, with variable party membership that changes frequently, species with more predictable subgroup composition also show mixed results. Yunnan snub-nosed monkeys (*Rhinopithecus bieti*) displayed an increase in fission events and subgrouping when important seasonal components of their diet were available, but not in relation to predator presence [50]. Another study on the same species found no effect of food availability on subgrouping [51]. In hamadryas baboons, bands fissioned more frequently during warmer and wetter months, coinciding with time periods during which preferred food items such as doum palm fruits (*Hyphaene* spp.) were less abundant – likely as a strategy to mitigate feeding competition during these months [13, 52, 53].

One reason for the lack of an effect of predator presence on the temporal cohesion between parties may have been the consistently high levels of predator presence throughout the study period so that the perceived risk for the baboons did not vary, even when we, according to our proxies, determined a lower predator presence. Although our dataset showed considerable variation in predator presence (Fig. 3), there were no periods entirely free of or particularly low in predator encounters. For instance, within any 2-day interval, there were at least two predator encounters. Such a constantly high predator presence may have prevented Guinea baboon parties from adjusting their spatial cohesion even if predator presence temporarily decreased slightly. Additionally, our measures of predator presence were perhaps too coarse to reliably capture changes in the perceived risk that might affect the grouping and space use of our study parties.

Our findings may also be in part affected by our relatively coarse assessment of food availability. We monitored the phenology of a proportion of known feeding tree species at a monthly interval. We did not, however, monitor all known feeding tree species or herbaceous plant foods consumed by our study parties, nor did we assess the distribution of feeding trees across the landscape. We were also unable to determine the abundance and distribution of other food types, such as insects readily consumed by our study parties.

Interestingly, in 2022, the period of lowest food availability according to our selection of feeding tree species was in June, July, and August, during the rainy season. Many grass shoots and herbaceous plant species became available during this time, and our study parties fed on them. Additionally, this period coincided with a mass occurrence of caterpillars of a species of hawk moths (Sphingidae), which made up 65% of our study parties’ feeding time in July. For comparison, in other months, insects made up only 4% of our study parties’ feeding time on average.

Therefore, our study parties could likely tolerate prolonged periods of low food availability. The lack of an effect of food availability, as we determined it, on the cohesion between parties may result from very low levels of feeding competition between parties in the study area so that even when fewer feeding trees are bearing fruit, alternative food sources are sufficiently available. The benefits of group living (e.g., support in conflicts, protection against predators) outweigh the costs related to feeding competition in our study population in Simenti, Senegal, thus allowing parties to stay in close spatial proximity year-round. However, it is possible that resource partitioning and competition are less relevant at the landscape level, but become more pronounced at a finer scale, such as within feeding patches or at the level of units rather than parties. At this more granular level, competition may be more immediate and intense, potentially leading to larger spacing between individuals foraging together, or higher levels of aggression in case of strong contest competition for highly valuable, contestable resources.

## Conclusion

We found no evidence for spatial or temporal avoidance between neighbouring Guinea baboon parties. On the contrary, for most of the observed parties, we found strong evidence for attraction-based space use patterns. Fission-fusion dynamics at the party level in our study period did not act as a strategy to mitigate resource competition or the risks associated with predator presence. Most study parties stayed in close spatial proximity to at least one other party, irrespective of variation in these ecological variables. The observed patterns indicate low levels of resource competition between parties when assessed over their entire home ranges. Whether this is a general feature of Guinea baboons or specific to our study parties and the ecology in Simenti, Senegal, remains to be investigated.

## Declarations

### Availability of data and materials

GPS data from 2010 to 2012 can be found on Göttingen Research Online: https://doi.org/10.25625/IHEZUE [21]. GPS and phenological data, as well as data on predator presence from 2022 are available from the corresponding author or from JF upon request.

### Competing interests

The authors declare that they have no competing interests.

### Funding

This research was supported by the Deutsche Forschungsgemeinschaft (DFG, German Research Foundation), Grant/Award Number: 254142454 / GRK 2070.

### Authors’ contributions

LO, JF and DZ designed the study. JB equipped Guinea baboon males with GPS collars and provided a description of the collaring procedure. LO collected the data in 2022. RM and LO analysed the data and prepared the figures. LO prepared the manuscript with contributions from all co-authors. All authors read, edited and approved the final manuscript.

## Acknowledgements

We are grateful to the Direction des Parcs Nationaux (DPN) and the Ministère de l’Environnement et de la Protéction de la Nature (MEPN) de la République du Sénégal for the approval to conduct this study in the Niokolo-Koba National Park. We especially thank the former conservateur of the park, Jacques Gomis, for his support. We are grateful to all the CRP Simenti staff and field assistants for their support in the field, in particular Djibril Coly, Chérif Younousse Kéba Camara, and Amadou Bamba Diedhiou. We owe special thanks to Irene Gutiérrez Díez for her help with the data collection in the field.

## Notes

https://doi.org/10.25625/IHEZUE

### Competing Interest Statement

The authors have declared no competing interest.

