## Supplementary material for "Patterns of landscape partitioning indicate low levels of resource competition among neighbouring Guinea baboon (*Papio papio*) parties"

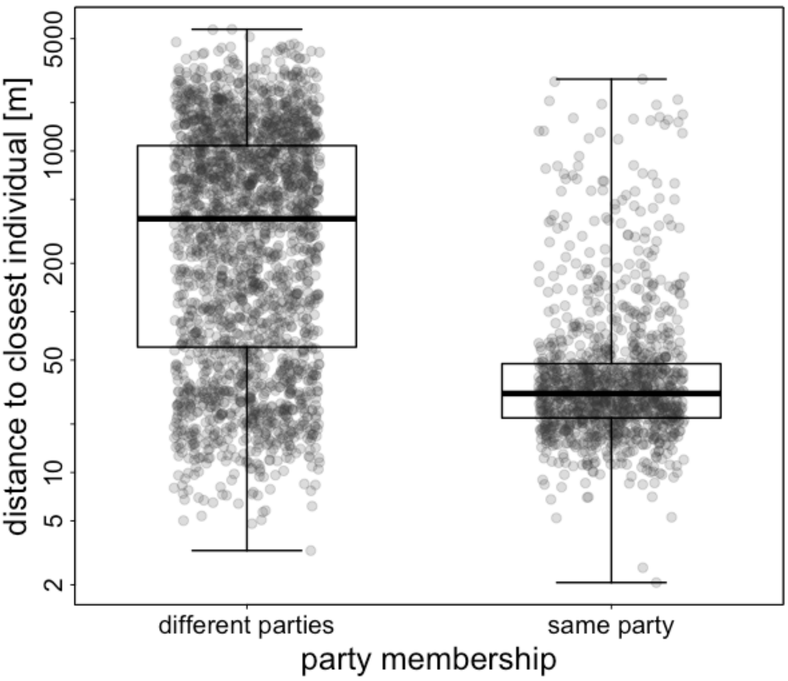


Fig. S1: Minimum distances between individuals of the same parties compared to individuals of different parties that were collared from 2010 to 2012. Boxplots depict the median (black line) and IQR with the lower (25%) and upper (75%) quartile. Whiskers represent the 2.5$^{th}$ and 97.5$^{th}$ percentiles. The distance to the closest party is depicted on a log-scale for visual clarity.

Table S1: Tree species subject to phenological monitoring and respective plant parts that baboons fed on more than once according to Klapproth et al. (unpublished data) (1 = occasional food resource, 2 = common food resource, 3 = key food resource).

| Family | Species | used | parts eaten |
|---|---|---|---|
| Anacardiaceae | *Lannea microcarpa* | 3 | fruits |
| Annonaceae | *Hexalobus monopetalus* | 2 | fruits, flowers |
| | *Spondias mombin* | 2 | fruits |
| Apocynaceae | *Saba senegalensis* | 1 | fruits |
| Arecaceae | *Borassus akeassii* | 3 | fruits, fibre, petiole |
| Bombacaceae | *Bombax costatum* | 2 | seeds |
| Caesalpinioideae | *Cordyla pinnata* | 2 | fruits |
| | *Piliostigma* spp. | 2 | pods, seeds |
| | *P. reticulatum* | | |
| | *P. thoningii* | | |
| | *Tamarindus indica* | 2 | fruits |
| Capparaceae | *Capparis tomentosa* | 1 | seeds |
| Combretaceae | *Combretum* spp. | 2 | seeds |
| | *C. glutinosum* | | |
| | *C. micranthum* | | |
| | *Terminalia macroptera* | 2 | seeds, bark |
| Ebenaceae | *Diospyros mespiliformes* | 1 | fruits |
| Loganiaceae | *Strychnos spinosa* | 2 | fruits |
| Mimosiodeae | *Mimosa pigra* | 2 | pods, seeds |
| | *Acacia* spp. | 2 | pods, seeds, flowers |
| | *A. macrostachya* | | |
| | *A. seyal* | | |
| | *A. sieberiana* | | |
| | *Dicrostachys cinera* | 2 | pods, seeds |
| | *Parkia biglobosa* | 2 | fruits |
| Moraceae | *Ficus ingens* | 2 | fruits |
| Papilionoideae | *Pterocarpus erinaceus* | 3 | seeds, bark |
| Rhamnaceae | *Ziziphus* spp. | 2 | fruits |
| | *Z. mauretania* | | |
| | *Z. mucronata* | | |
| Rubiaceae | *Sarcocephalus latifolius* | 2 | fruits |
| Tiliaceae | *Grewia* spp. | 2 | fruits |
| | *Grewia bicolor* | | |
| | *Grewia lasiodiscus* | | |
| Ulmaceae | *Celtis integrifolia* | 2 | fruits |
| Verbenaceae | *Vitex madiensis* | 1 | fruits |

Table S2: Overlap between home ranges (KDE, 95% contour level) of parties observed in 2010. Values represent the proportion of overlap (0 = no overlap, 1= full overlap). The extent of the overlap is indicated as a colour gradient from green (low overlap) to yellow (high overlap).

| | P4 | P5 | P6 | P9 |
|---|---|---|---|---|
| P4 | | 0.68 | 0.72 | 0.92 |
| P5 | 0.95 | | 1.00 | 0.94 |
| P6 | 0.93 | 0.92 | | 0.92 |
| P9 | 0.82 | 0.64 | 0.69 | |

Table S3: Overlap between home ranges (KDE, 95% contour level) of parties observed from 2011 to 2012. Values represent the proportion of overlap (0 = no overlap, 1= full overlap). The extent of the overlap is indicated as a colour gradient from green (low overlap) to yellow (high overlap).

| | P4 | P5 | P6 | P9 |
|---|---|---|---|---|
| P4 | | 0.83 | 0.81 | 0.81 |
| P5 | 0.88 | | 0.96 | 0.73 |
| P6 | 0.92 | 0.96 | | 0.82 |
| P9 | 0.95 | 0.80 | 0.77 | |

Table S4: Overlap between core areas (KDE, 50% contour level) of parties observed in 2010. Values represent the proportion of overlap (0 = no overlap, 1= full overlap). The extent of the overlap is indicated as a colour gradient from green (low overlap) to yellow (high overlap).

| | P4 | P5 | P6 | P9 |
|---|---|---|---|---|
| P4 | | 0.71 | 0.80 | 0.99 |
| P5 | 0.63 | | 0.98 | 0.65 |
| P6 | 0.59 | 0.86 | | 0.61 |
| P9 | 0.85 | 0.67 | 0.76 | |

Table S5: Overlap between core areas (KDE, 50% contour level) of parties observed from 2011 and 2012. Values represent the proportion of overlap (0 = no overlap, 1= full overlap). The extent of the overlap is indicated as a colour gradient from green (low overlap) to yellow (high overlap).

| | P4 | P5 | P6 | P9 |
|---|---|---|---|---|
| P4 | | 0.80 | 0.72 | 0.87 |
| P5 | 0.51 | | 0.97 | 0.61 |
| P6 | 0.56 | 0.93 | | 0.59 |
| P9 | 0.88 | 0.83 | 0.68 | |

Table S6: Model results on spatiotemporal landscape partitioning between neighbouring parties in response to food availability and number of predator encounters within 7 days (estimates, standard errors, 95% credible intervals, Rhat, as well as Bulk and Tail Effective Sample Sizes).

| Term | Estimate | Est.Error | $CI_{lower}$ | $CI_{upper}$ | Rhat | Bulk_ESS | Tail_ESS |
|---|---|---|---|---|---|---|---|
| Intercept | 5.15 | 0.71 | 3.67 | 6.49 | 1.00 | 1221 | 1826 |
| pred.enc.7 | -0.00 | 0.01 | -0.02 | 0.02 | 1.00 | 2909 | 2878 |
| food.score | 0.19 | 0.52 | -0.80 | 1.30 | 1.00 | 2755 | 2684 |

Table S7: Model results on spatiotemporal landscape partitioning between neighbouring parties in response to food availability and number of predator encounters within 30 days (estimates, standard errors, 95% credible intervals, Rhat, as well as Bulk and Tail Effective Sample Sizes).

| Term | Estimate | Est.Error | $CI_{lower}$ | $CI_{upper}$ | Rhat | Bulk_ESS | Tail_ESS |
|---|---|---|---|---|---|---|---|
| Intercept | 5.49 | 0.84 | 3.79 | 7.14 | 1.00 | 975 | 1470 |
| pred.enc.30 | -0.01 | 0.01 | -0.02 | 0.01 | 1.00 | 1555 | 1966 |
| food.score | 0.35 | 0.53 | -0.64 | 1.43 | 1.00 | 1866 | 1944 |

Table S8: Model results on spatiotemporal landscape partitioning between neighbouring parties in response to food availability and number of predator encounters within two days (estimates, standard errors, 95% credible intervals, Rhat, as well as Bulk and Tail Effective Sample Sizes).

| Term | Estimate | Est.Error | $CI_{lower}$ | $CI_{upper}$ | Rhat | Bulk_ESS | Tail_ESS |
|---|---|---|---|---|---|---|---|
| Intercept | 5.00 | 0.63 | 3.63 | 6.19 | 1.01 | 731 | 925 |
| pred.enc.2 | 0.03 | 0.03 | 0.03 | 0.08 | 1.00 | 1391 | 1667 |
| food.score | 0.13 | 0.54 | 0.54 | 1.24 | 1.00 | 1589 | 1797 |

Table S9: Model results on spatiotemporal landscape partitioning between neighbouring parties **at noon** in response to food availability and number of predator encounters within two weeks (estimates, standard errors, 95% credible intervals, Rhat, as well as Bulk and Tail Effective Sample Sizes).

| Term | Estimate | Est.Error | $CI_{lower}$ | $CI_{upper}$ | Rhat | Bulk_ESS | Tail_ESS |
|---|---|---|---|---|---|---|---|
| Intercept | 4.36 | 0.72 | 2.84 | 5.79 | 1.00 | 1342 | 1948 |
| pred.enc.14 | 0.01 | 0.01 | -0.01 | 0.04 | 1.00 | 3795 | 2875 |
| food.score | -0.30 | 0.70 | -1.66 | 1.08 | 1.00 | 3370 | 2584 |

Table S10: Model results on spatiotemporal landscape partitioning between neighbouring parties **at noon** in response to food availability and number of predator encounters within 7 days (estimates, standard errors, 95% credible intervals, Rhat, as well as Bulk and Tail Effective Sample Sizes).

| Term | Estimate | Est.Error | $CI_{lower}$ | $CI_{upper}$ | Rhat | Bulk_ESS | Tail_ESS |
|---|---|---|---|---|---|---|---|
| Intercept | 4.37 | 0.74 | 2.89 | 5.80 | 1.00 | 1161 | 1705 |
| pred.enc.7 | 0.01 | 0.01 | -0.01 | 0.04 | 1.00 | 4081 | 2944 |
| food.score | -0.31 | 0.66 | -1.69 | 0.94 | 1.00 | 3747 | 3000 |

Table S11: Model results on spatiotemporal landscape partitioning between neighbouring parties **at noon** in response to food availability and number of predator encounters within 30 days (estimates, standard errors, 95% credible intervals, Rhat, as well as Bulk and Tail Effective Sample Sizes).

| Term | Estimate | Est.Error | $CI_{lower}$ | $CI_{upper}$ | Rhat | Bulk_ESS | Tail_ESS |
|---|---|---|---|---|---|---|---|
| Intercept | 4.68 | 0.84 | 3.01 | 6.37 | 1.00 | 1310 | 2200 |
| pred.enc.30 | -0.00 | 0.01 | -0.02 | 0.01 | 1.00 | 2306 | 2326 |
| food.score | -0.03 | 0.81 | -1.70 | 1.58 | 1.00 | 2594 | 2165 |

Table S12: Model results on spatiotemporal landscape partitioning between neighbouring parties **at noon** in response to food availability and number of predator encounters within 2 days (estimates, standard errors, 95% credible intervals, Rhat, as well as Bulk and Tail Effective Sample Sizes).

| Term | Estimate | Est.Error | $CI_{lower}$ | $CI_{upper}$ | Rhat | Bulk_ESS | Tail_ESS |
|---|---|---|---|---|---|---|---|
| Intercept | 4.46 | 0.68 | 3.06 | 5.86 | 1.01 | 1291 | 1974 |
| pred.enc.2 | 0.04 | 0.03 | -0.02 | 0.09 | 1.00 | 3587 | 2268 |
| food.score | -0.25 | 0.69 | -1.60 | 1.13 | 1.00 | 3295 | 2414 |
